# Intracellular and Dual-Site Inhibition of a Bitter Taste GPCR

**DOI:** 10.64898/2025.12.04.692259

**Authors:** Nitsan Dallal, Gil Daniel Paz, Noga Nir Marom, Yael Keselman, Shir Eyal, Evgenii Ziaikin, Alon Rainish, Lior Peri, Einav Malach, Masha Y. Niv

## Abstract

Bitter taste receptors (TAS2Rs) are G protein-coupled receptors expressed in both gustatory and extraoral tissues and activated by a broad range of compounds. TAS2R14 is among the most promiscuous members of this family, responding to many structurally diverse ligands. Cryo-electron microscopy structures of TAS2R14 have revealed agonists binding in an intracellular pocket, raising the question of the main sites of interaction for known TAS2R14 antagonists. To address this, we examined the effects of mutations at residues located in the extracellular and intracellular regions on receptor inhibition by three antagonist compounds. Mutations in the extracellular region reduced the inhibitory effect of LF22, whereas all three compounds showed reduced inhibition in the intracellular mutants. Computational co-folding of these ligands with TAS2R14 supported these observations, indicating that LF22 interacts with both top and bottom binding sites, whereas LF1 and probenecid engage predominantly the intracellular site adjacent to the G protein interface. Interestingly, LF1 is much more potent for TAS2R16 than its known inhibitor probenecid. These findings reveal distinct inhibitory mechanisms among TAS2R antagonists and provide new insights towards designing inhibitors of bitter taste.

## 1. Introduction

G protein-coupled receptors (GPCRs) represent the largest family of cell-surface receptors in humans, with nearly 800 members involved in transducing extracellular signals into cellular responses^1^. They regulate numerous physiological processes and are targeted by over one-third of all FDA-approved drugs, of which more than half act as antagonists^1^. Within this family, bitter taste receptors (TAS2Rs) form a small but distinctive subfamily of 25 functional receptors in humans^2–6^. Human TAS2Rs detect over two thousand structurally diverse natural and synthetic bitter molecules and play a crucial role in taste perception^7^. In addition, these receptors are implicated in extra-oral functions such as immune defense, airway smooth muscle relaxation, and cancer biology^8–10^.

Among TAS2Rs, TAS2R14 is the most promiscuous member, activated by structurally diverse ligands, including flavonoids, alkaloids and peptides^7,11^. TAS2R14 is expressed in taste buds and in multiple extra-oral tissues, such as the respiratory tract, placenta, thyroid, and pancreas, where it has been implicated in both protective and pathological processes^12–14^. Consistent with this dual role, high TAS2R14 expression has been linked to improved survival in pancreatic ductal adenocarcinoma, whereas in other cancers it correlates with poorer prognosis^12,15^. This broad expression pattern and ligand diversity highlights TAS2R14 as a receptor of major physiological and therapeutic relevance^4^.

Contrary to the wealth of agonists identified, only a few TAS2R14 antagonists have been described to date, i.e., ^16,17^. For example, our virtual screening discovery efforts yielded a small set of TAS2R14 antagonists, which were later validated as chemical probes and shown to inhibit TAS2R14 activity in several cancer cell lines^13,16^. Specifically, LF1 and LF22 compounds (Figure 1) were found based on their fit to the canonical extracellular binding pocket of the three-dimensional model of the TAS2R14 receptor, traditionally assumed to accommodate both agonists and antagonists^11,16,18^. After the discovery of these antagonists, several cryo-electron microscopy (Cryo-EM) structures of TAS2R14 were solved, providing unprecedented insight into its architecture^19–22^. The experimental structures unexpectedly revealed an additional intracellular agonist binding pocket. In other GPCR families, intracellular binding pockets have been detected in a few cases and usually associated with antagonist binding^23–28^. This raised the possibility that TAS2Rs might also be inhibited through an intracellular site. However, no experimental TAS2R14 structure has been solved with an antagonist, leaving open the question of where antagonists interact with this receptor.

**Figure 1.**
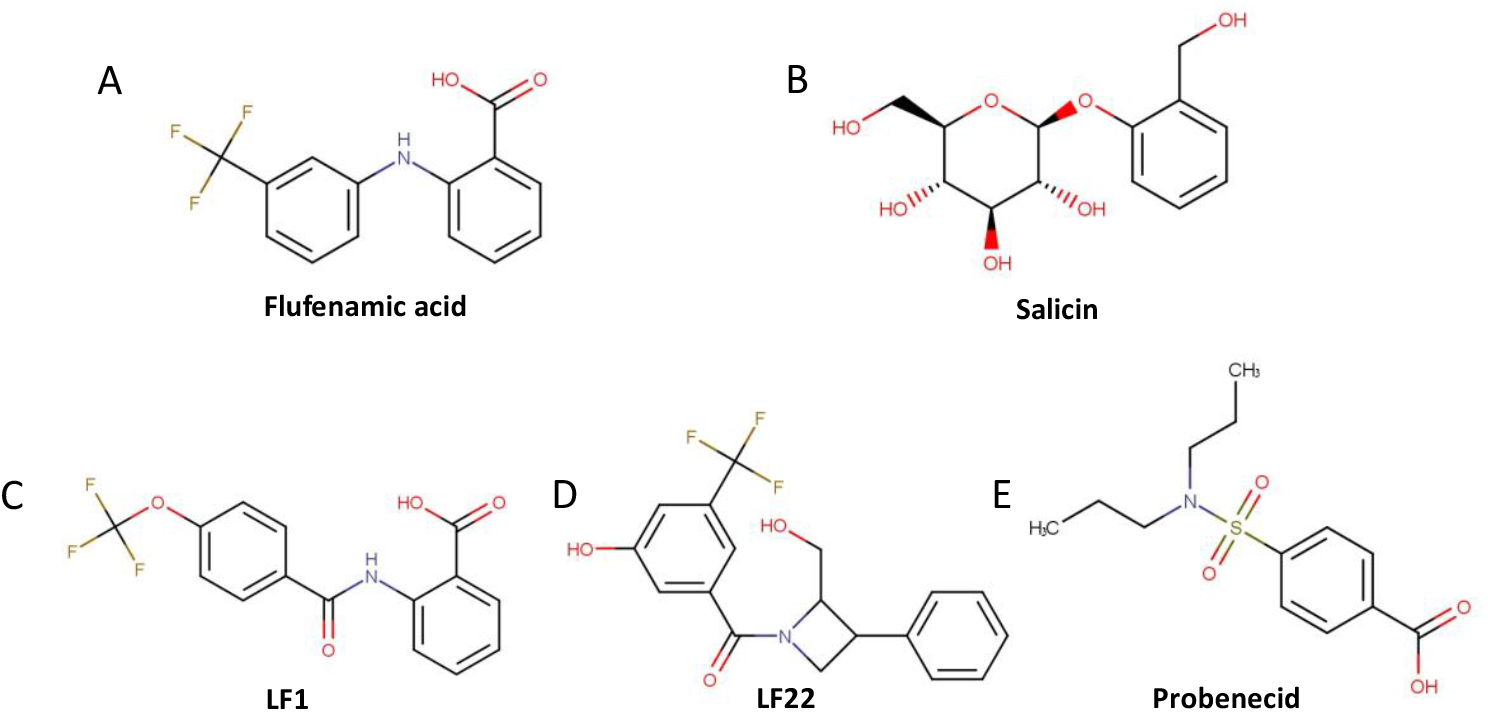
Agonist and antagonist used in this research.

In this work, we explored whether TAS2R14 antagonists LF1 and LF22 act through the extracellular pocket (for which they were originally designed), the intracellular pocket, or both. We also included probenecid, known to inhibit TAS2R16^29^, and recently shown to inhibit also TAS2R14^30^, and tested all three compounds on TAS2R16 to provide a comparative reference. Using functional assays of representative mutants, we characterized the binding locations of these antagonists and provided computational models compatible with the experimental data. Together, this approach expands mechanistic insights into bitter receptors inhibition and lays the groundwork for rational design of targeted bitter blockers.

## 2. Materials and methods

### Cell culture

Experiments were performed using HEK293T cells. Cells were cultured in plates with 10% Dulbecco’s Modified Eagle’s Medium (DMEM) containing 10% Fetal Bovine Serum (FBS), 1% l-glutamine amino acid, and 0.2% penicillin streptomycin. Every 3-4 days, when plates were at 80% confluence, the medium was removed, and cells were transferred into a fresh growth medium. Cells were kept in the incubator, in 37 °C and 5% CO_2_.

### IP-One assay

For functional expression of the human bitter taste receptor TAS2R14 or TAS2R16, HEK293T cells were grown at about 80% confluence and transiently transfected with 2 μg of a plasmid (pcDNA3.1) encoding N-terminally modified TAS2R14 or TAS2R16 (N-terminal addition of a cleavable HA-signal peptide, followed by a FLAG-tag and the first 45 amino acids of the rat somatostatin receptor 3) and 1 μg of a plasmid encoding the chimeric Gαqi5 protein, as described in previous work^16^. Transfections were performed using Mirus TransIT-293 (Mirus Bio) at a 1:3 DNA to reagent ratio.

24 h after transfection, the transfected cells were suspended in 10% DMEM and seeded onto a 384-well culture micro plate (Greiner) at a density of 5×10^5^ (cells/ml). The plate was then kept in an incubator for 24 h to obtain cell adherence. The next day, cell culture medium supernatant was removed from the plates and stimulation buffer was added to each well. Cells were then treated by the addition of the tested compounds. Plates were incubated for 150 min, to allow IP1 accumulation inside the cell^16,31^. For investigation of TAS2R14 or TAS2R16 inhibition, the cells were first pre-incubated for 30 min with potential inhibitor, followed by addition of either 1 µM FFA or 10 mM salicin and continuing incubation for further 150 min.

Accumulation of second messenger was stopped by adding detection reagents (IP1-d2 conjugate and Anti-IP1cryptate TB conjugate) dissolved in kit lysis buffer (HTRF IP-One Gq Detection Kit, Revvity). FRET-ratios were calculated as the ratio of emission intensity of the FRET acceptor (665/10 nm) divided by the FRET donor intensity (620/10 nm) (ClarioStar, BMG). Raw FRET-ratios were normalized to buffer conditions (0%) and the maximum effect of FFA or Salicin (100%), and the obtained responses were analyzed using the equation for sigmoid concentration-response curves (four-parameter) implemented in GraphPad Prism for Windows (GraphPad Software, La Jolla, USA) to derive the maximum effect (E_max_, relative to Salicin) and the ligand potency (EC_50_ or IC_50_). For antagonist properties, the maximum effect at 1 µM FFA or 10 mM salicin was normalized to 100%.

### Compounds

Reagents and compounds were obtained from the following suppliers: Thermo Fisher Scientific (USA), Sigma (USA; flufenamic acid, salicin, probenecid), Gibco (USA; DMEM, FBS), Revvity (France; IP-One HTRF kit), BLDpharm (China), Enamine (Ukraine; LF1 (Cat. No. Z85879385) and LF22 (Cat. No. Z3906872218)), and Mirus Bio (USA; TransIT-293 reagent). All Compounds were dissolved in DMSO. Serial dilutions of compounds were prepared in the stimulation buffer of the kit (containing 50 mM LiCl to prevent IP1 degradation) at the desired working concentration on the day of the experiment (Cisbio IP-ONE-Gq KIT).

### Co-folding of protein-ligand complexes

Boltz-2 was used to predict the structure of the protein-ligand complex using the default mmseqs2 server to generate MSA, with a diffusion sample number of 20^32,33^. The input data consisted of ligand SMILES taken, where possible, from PubChem^34^ or from the original publications in which LF1 and LF22 were discovered^16^. The TAS2R sequences were taken from BitterDB^7^.

## 3. Results

Recent cryo-EM structures of TAS2R14, solved independently by different groups, consistently revealed two binding sites within the transmembrane core: an extracellular (top, canonical) pocket and an intracellular (bottom) pocket at the receptor-G protein interface^19–22^. In PDB 8RQL, two copies of flufenamic acid (FFA) simultaneously occupy both binding sites, an uncommon scenario featuring two copies of the same agonist in separate pockets^20^ (Figure 2A-C).

**Figure 2.**
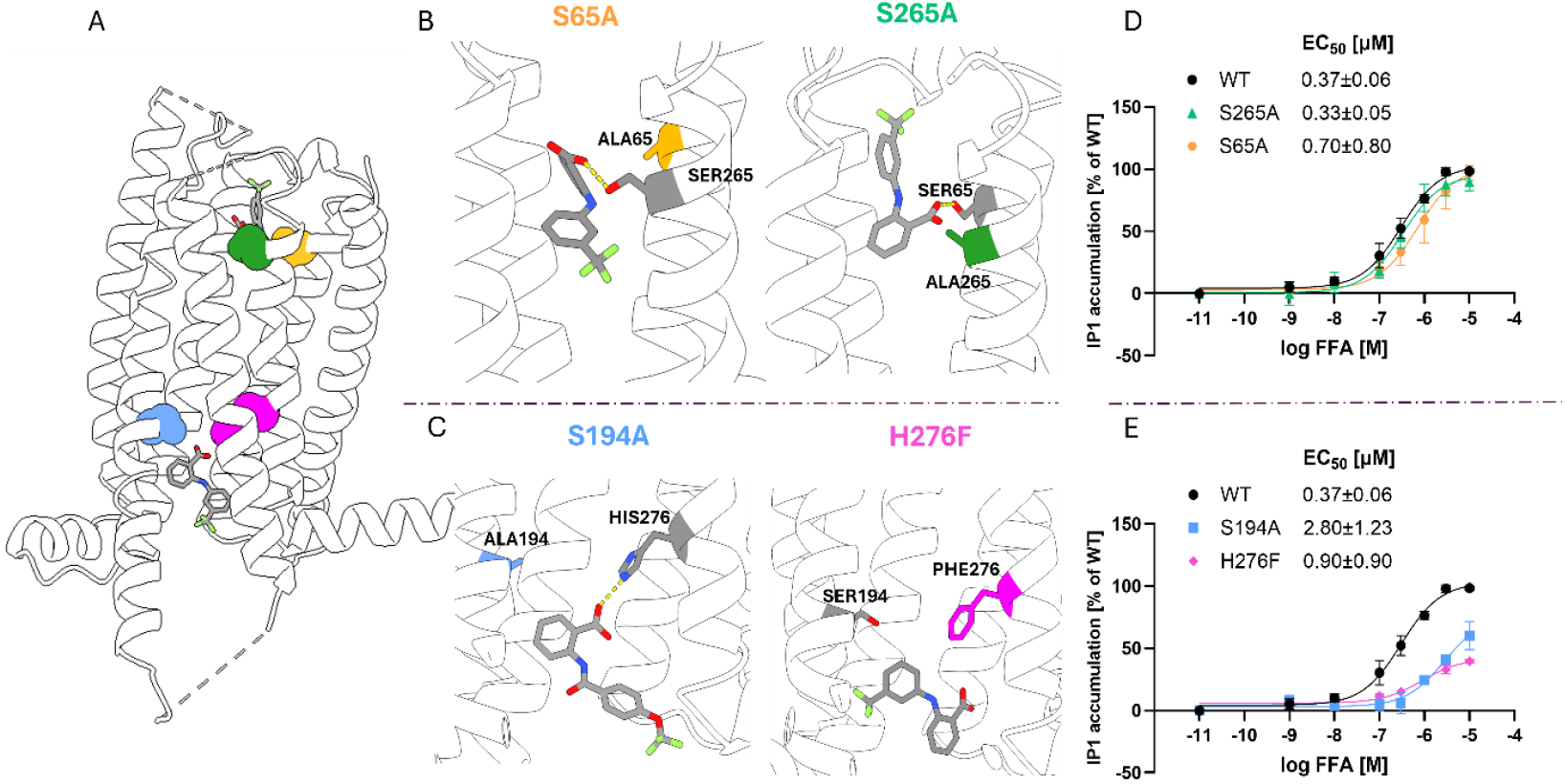
Binding sites and mutations in TAS2R14. Effect of upper and lower site mutations on TAS2R14 activation by FFA. (A) Overall Cryo-EM structure of TAS2R14 with FFA bound in both the upper and lower pockets, with mutated residues highlighted (S65^2.60^ in orange, S265^7.38^ in green, S194^5.54^ in light blue and H276^7.49^ in magenta). Hydrogen bonds are shown in yellow. Prepared with ChimeraX^39^. (B) Close up view of the upper binding site showing FFA and mutation in S65^2.60^ or S265^7.38^. (C) Close up view of the lower binding site showing FFA and mutation in S194^5.54^ or H276^7.49^. (D-E) HEK293T cells were transfected with either TAS2R14 WT or mutants, and with Gαqi5, and IP-One accumulation was measured. (D) Dose response curves of WT and upper site mutants (S65A, S265A). (E) Dose response curves of WT and lower site mutants (S194A, H276F).^43^

To determine where known antagonists interact with the receptor, we evaluated how mutations at key residues influence FFA-induced activation. We focused on positions previously highlighted by structural and mutagenesis studies as important for either the top or the bottom pockets. In the top pocket, we examined S65A^2.60^ and S265A^7.38^ (positions annotation follows the Ballesteros-Weinstein scheme for GPCRs^35^). Residue S65^2.60^ forms polar interactions with the amine linker of FFA, and substitutions at this position influence agonist recognition^36^. S265^7.38^ corresponds to a position where naturally occurring variants have been identified and has been reported to affect agonist potency and efficacy^11,20,37^. In the bottom pocket, we tested S194A^5.54^ and H276F^7.49^. Both residues directly interact with FFA in the Cryo-EM structure of TAS2R14^20^, and previous work has shown that both mutations impair receptor activation^20^. Position 7.49 is a highly conserved histidine present in 24 of the human TAS2Rs^20,38^. We measured FFA-induced responses in HEK293T cells expressing either wild-type (WT) and mutant receptors (Figure 2D-E).

WT exhibited an EC_50_ of 0.37 ± 0.06 µM, in agreement with previous studies^20,21^. In the top binding site, both S65A^2.60^ (EC_50_ = 0.70 ± 0.08 µM) and S265A^7.38^ (EC_50_ = 0.33 ± 0.05 µM) showed EC_50_ values similar to WT and both mutants reached maximal activation of approximately 100% (Figure 2D). In the bottom binding site, S194A^5.54^ displayed a marked decrease in potency (EC_50_ = 2.80 ± 1.23 µM) together with a reduced maximal activation to about 60%. H276F^7.49^ also showed a rightward shift (EC_50_ = 0.90 ± 0.09 µM) and an even stronger reduction in maximal activation to approximately 40% (Figure 2E). Together, these results indicate that mutations in the bottom pocket have a clear functional impact on FFA activation, whereas the effects of the top-pocket mutations are minimal. Importantly, all the mutants still respond to FFA in a dose-dependent manner, allowing the next step, namely testing the effects of antagonists.

### LF1, LF22 and probenecid

We next asked how antagonists interact with TAS2R14. LF1 and LF22 were originally found through predicted compatibility with the canonical binding site of TAS2R14, prior to the availability of the TAS2R14 cryo-EM structures and before the identification of a second intracellular pocket^16^. In addition to LF1 and LF22, we tested probenecid, a small-molecule drug previously characterized as a TAS2R16 antagonist, suggested to act via the TAS2R16 intracellular site^29,40^ and recently shown to inhibit TAS2R14^30^.

In WT, LF1 inhibited TAS2R14 with an IC_50_ of 18.0 ± 0.50 µM and reduced responses to 44% of FFA activity. In the top site mutants, S65A^2.60^ and S265A^7.38^ displayed IC_50_ values of 3.0 ± 8.10 µM and 11.6 ± 1.70 µM with inhibition levels of 47% and 55%, respectively (Figure 3A). In the bottom site mutants, S194A^5.54^ showed an IC_50_ of 2.90 ± 1.40 µM and 59% inhibition, while H276F^7.49^ completely abolished LF1 inhibition (Figure 3B). Taken together, top pocket mutations had minimal influence on LF1 inhibition, while bottom pocket mutations, particularly H276F^7.49^, strongly disrupted LF1 activity.

**Figure 3.**
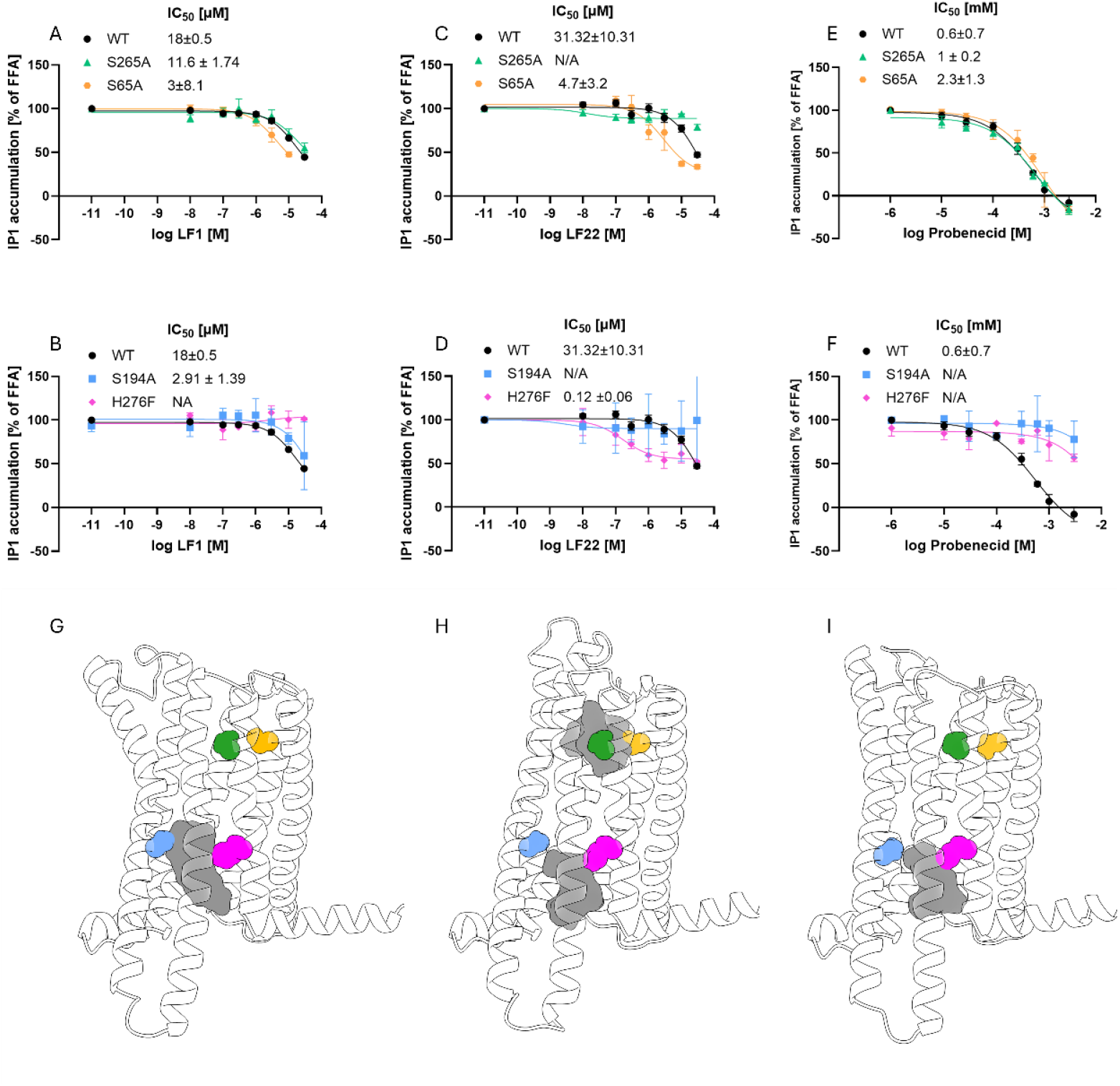
Inhibition of TAS2R14 WT and mutants. Dose response curves showing inhibition of TAS2R14 by LF1, LF22, and probenecid and predicted antagonist binding poses for TAS2R14. Panels A to F show IP-One accumulation in HEK293T cells transfected with TAS2R14 WT or mutants S65A, S265A, S194A, and H276F together with Gαqi5, measured in the presence of 1 µM FFA (EC_80_) and increasing concentrations of antagonists. (A) Inhibition of FFA responses by LF1 in WT (IC_50_ = 18.0 ± 0.50 µM), S265A (IC_50_ = 11.6 ± 1.74 µM), and S65A (IC_50_ = 30 ± 8.10 µM). (B) Inhibition of FFA responses by LF1 in WT (IC_50_ = 18 ± 0.50 µM), S194A (IC_50_ = 2.91 ± 1.39 µM), and H276F (no inhibition). (C) Inhibition of FFA responses by LF22 in WT (IC_50_ = 31.32 ± 10.31 µM), S265A (no inhibition), and S65A (IC_50_ = 4.70±3.20 µM). (D) Inhibition of FFA responses by LF22 in WT (IC_50_ = 31.32 ± 10.31 µM), S194A (no inhibition), and H276F (IC_50_ = 0.12±0.06 µM). (E) Inhibition of FFA responses by probenecid in WT (IC_50_ = 0.60 ± 0.70 mM), S265A (IC_50_ = 1 ± 0.20 mM) and S65A (IC_50_ = 2.30 ± 1.30 mM). (F) Inhibition of FFA responses by probenecid in WT (IC_50_ = 0.60 ± 0.70 mM), while both S194A and H276F showed no inhibition. All data are presented as mean ± SEM of 3-6 replicates. (G-I). Boltz-2 predicted binding pose ensemble density surfaces of LF1 (G), LF22 (H), probenecid (I), based on five independent co-folding simulations for each ligand. Residues are highlighted: S65^2.60^ in orange, S265^7.38^ in green, S194^5.54^ in light blue, and H276^7.49^ in magenta.

LF22 inhibited FFA-induced responses in WT TAS2R14 with an IC_50_ of 31.30 ± 10.30 µM and reduced responses to 47%. In the top-site mutants, S65A^2.60^ displayed an IC_50_ of 4.70 ± 3.20 µM with 33% inhibition, while S265A^7.38^ showed no measurable inhibition (Figure 3C). In the bottom site mutants, S194A^5.54^ abolished LF22 inhibition, whereas H276F^7.49^ increased potency with an IC_50_ of 0.12 ± 0.06 and inhibition level of 52% (Figure 3D). These results show that LF22 inhibition is disrupted by mutations in both pockets, specifically by the top pocket mutant S265A^7.38^ and the bottom pocket mutant S194A^5.54^.

Probenecid completely abolished the WT response to FFA, with an IC_50_ of 0.60 ± 0.70 mM. In the top site mutants S65A^2.60^ and S265A^7.38^, probenecid retained its full inhibitory capacity, showing IC_50_ values of 2.30 ± 1.30 mM and 1.0 ± 0.20 mM, respectively (Figure 3E). In contrast, mutations in the bottom binding pocket eliminated inhibition: both S194A^5.54^ and H276F^7.49^ failed to respond to probenecid and showed no reduction in FFA-induced activity (Figure 3F).

To visualize possible mode of interactions of antagonists with TAS2R14, we used Boltz-2, a structure-based deep-learning framework that integrates protein flexibility and ligand sampling to predict binding poses^32^. A key feature of Boltz-2 is its ability to model ligand binding without predefining a binding pocket, enabling unbiased exploration of possible interaction sites across the receptor surface. Boltz-2 takes the receptor sequence as input and jointly samples receptor conformations and protein ligand interactions^32^. For each antagonist, five independent simulations were performed, and the resulting poses were clustered to identify the most probable binding modes.

Across the five predicted poses, LF1 was consistently placed in the bottom pocket (Figure 3G). LF22 produced a mixed pattern, with two poses positioned in the bottom pocket and three located in the top pocket (Figure 3H). Probenecid localized to the intracellular site in all predicted complexes (Figure 3I).

### Antagonists cross-reactivity

Probenecid is one of the few established TAS2R antagonists^29^. Mutations in the intracellular region reduced probenecid efficacy without affecting agonist activation, and probenecid lowered the maximal response to salicin without shifting its EC_50_, indicating that it decreases receptor efficacy rather than competing with the agonist at its binding site. The recent Cryo-EM structure of TAS2R16, solved with salicin bound in the extracellular binding site^41^, is compatible with this mechanism. Our results indicate that probenecid inhibits TAS2R14 through dependence on the intracellular site, and we next asked whether TAS2R14 antagonists also inhibit TAS2R16.

To examine TAS2R16 activation, we first measured the response to salicin in both WT and the N96T^3.43^ mutant, which lies within the intracellular region of the receptor and has been examined in previous studies^35^. In WT, salicin induced a clear dose-dependent activation with EC_50_ of 13.00 ± 4.30 mM. The N96T^3.43^ mutant showed a markedly stronger response, reaching 372% of the WT maximal activation, consistent with increased receptor sensitivity. This was also accompanied by a lower EC_50_ of 5.52 ± 0.62 mM (Figure 4A).

**Figure 4.**
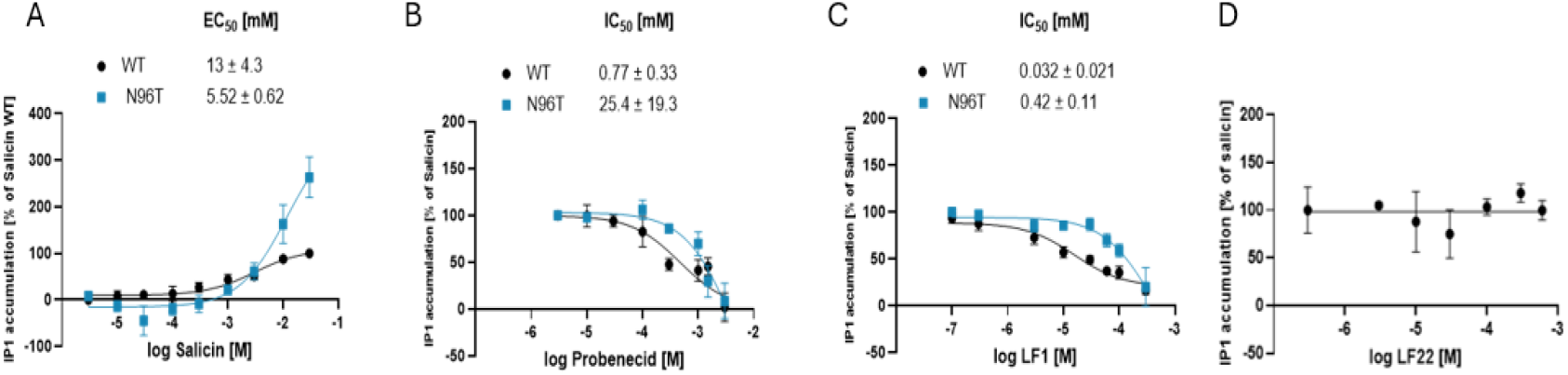
TAS2R16 activation and inhibition. HEK293T cells were transfected with TAS2R16 WT or N96T^3.43^, and Gαqi5-mediated IP-One accumulation was measured. (A) Salicin activated TAS2R16 in a dose-dependent manner with an EC_50_ of 13 ± 4.30 mM for WT and 5.52 ± 0.62 mM for N96T^3.43^. (B) Probenecid inhibited TAS2R16 activation induced by 10 mM salicin, with an IC_50_ of 0.77 ± 0.33 mM for WT, and 25.40 ± 19.30 mM for N96T^3.43^. (C) LF1 inhibited TAS2R16 activation induced by 10 mM salicin, with an IC_50_ of 0.032 ± 0.021 mM for WT and 0.42 ± 0.11 mM for N96T^3.43^. (D) LF22 did not inhibit TAS2R16 activation. Data represent mean ± SEM of 3-6 replicates, except for panel D where only one biological replicate was performed.

We next examined inhibition by probenecid. In WT, probenecid completely blocked salicin-induced activation, with an IC_50_ of 0.77 ± 0.33 mM. In N96T^3.43^, inhibition was substantially weaker, and the IC_50_ shifted to 25.40 ± 19.30 mM (Figure 4B).

LF1 also inhibited TAS2R16 in both WT and N96T^3.43^, reducing salicin-induced responses with IC_50_ values of 0.032 ± 0.021 mM and 0.42 ± 0.11 mM, respectively (Figure 4C). This is similar to probenecid effect in terms of compatability with intracellular inhibition, but the IC50 of LF1 is an order of magnitude lower than of the established TAS2R16 inhibitor probenecid. In contrast, LF22 did not inhibit salicin-induced activation of TAS2R16 (Figure 4D).

## 4. Discussion

Our study was motivated by our recent Cryo-EM finding of an intracellular binding site in TAS2R14^20^. Although 22 structures of TAS2R receptors have been solved to date, none have been determined in complex with antagonists. As a result, the structural basis for bitter taste receptor-antagonist interactions remains largely unknown. By combining mutagenesis with functional assays, we examined how alterations in the upper and lower binding pockets affect antagonist activity.

Our results show that LF1 and probenecid depend primarily on the intracellular pocket for inhibition. Both antagonists inhibited WT TAS2R14, and this inhibition persisted in the upper-site mutants S65A^2.60^ and S265A^7.38^, indicating that the extracellular pocket is not essential for their activity. In contrast, mutations in the lower pocket produced ligand-specific effects. The S194A^5.54^ mutation abolished inhibition by probenecid yet did not impair LF1 inhibition, which remained comparable to or slightly stronger than in the wild type. In contrast, H276F^7.49^ eliminated inhibition by both antagonists, demonstrating that this intracellular position is essential for their action. LF22 displayed a different pattern. It inhibited WT TAS2R14, yet its activity was lost in both S265A^7.38^ (upper-site) and S194A^5.54^ (lower-site) mutants, suggesting that LF22 relies on interactions in both pockets.

Boltz-2 co-folding simulations provided plausible 3D models that agree with these findings. Unlike many structure-based methods that require defining a binding region in advance, Boltz-2 determines ligand placement without any prior pocket specification. Across five simulations per ligand, LF1 and probenecid were repeatedly positioned in the intracellular pocket of TAS2R14, while LF22 yielded both top and bottom binding site poses. Together, these results reveal distinct antagonist classes, with LF1 and probenecid acting mainly through the intracellular pocket while LF22 depends on both pockets.

Because probenecid inhibits both TAS2R14 and TAS2R16, we asked whether this cross-receptor inhibition extends to additional antagonists. We examined probenecid, LF1 and LF22 effects on TAS2R16 activation by salicin. Probenecid and LF1 inhibited TAS2R16, and both showed reduced activity in the intracellular N96T^3.43^ mutant, mirroring their dependence on intracellular interactions in TAS2R14. LF22, which was the only one out of tested antagonists that was sensitive to top site mutations in TAS2R14, did not inhibit TAS2R16.

Together, our findings show that the more potent LF1 and the weaker probenecid inhibit both TAS2R14 and TAS2R16. For LF1, the IC_50_ values were 18.0 ± 0.50 µM for TAS2R14 and 32 ± 21 µM for TAS2R16, indicating comparable inhibitory potency for the two receptors, with slight preference for TAS2R14. Probenecid inhibited TAS2R14 with an IC_50_ of 0.60 ± 0.70 mM, and TAS2R16 with 0.77 ± 0.33 mM.

More broadly, our work highlights that TAS2Rs can host antagonists at intracellular sites. Antagonists may rely mainly on the bottom pocket or require combined top and bottom interactions. This may have implications for designing compounds with either wide or receptor-specific activity. ^5,44,45^ Our findings highlight the notion that intracellular binding sites should be taken into consideration. The variability we observed across the antagonists tested may be relevant for TAS2Rs in therapeutic contexts, where differences in receptor tuning and ligand behavior can guide the selection and design of compounds intended to target bitter receptors.

## Acknowledgements

We thank Asa Tirosh for help with mutagenesis and Michael Naim for fruitful discussions. Funding by ISF grants 1129/19 and 1096/25 is gratefully acknowledged.

## Authors contributions

ND: Functional and recruitment assays, formal analysis, visualization, writing and editing. GDP: Functional assays and original draft. NNM: Functional assays. YK: Functional assays. SE: Functional assays, review and editing. EZ and AR: Computational analyses and and visualization. LP: Resources, review and editing. EM: Supervision and project administration. MYN: Conceptualization, supervision, resources, writing and editing.

